# High-Density Microdroplet Cultivation Reveals the Essential Role of Microbial Interactions in the Growth of Environmental Microbes

**DOI:** 10.1101/2025.08.12.669842

**Authors:** Eun-Young Seo, Rikuta Suzuki, Yuki Takagi, Tomonori Kindaichi, Akiyoshi Ohashi, Setsu Kato, Yutaka Nakashimada, Yoshiteru Aoi

## Abstract

The significance of microbial interactions in nature is not well understood because of the limitations of techniques to assess microbial growth at the single-cell level under conditions of high cell density (more than 10^7^ cells/mL). To address this limitation for further solution of microbial uncultivability, we developed a Gel Microdroplet Aggregate in Oil (GMD-agg) cultivation method. This method provides a high inoculum cell density (>10^8^ cells/mL) but maintains pure cultures of each growth unit. Millions of hydrogel particles (10–30 µm in diameter) entrapping single cells with medium softly aggregated in oil (GMD-oil cultivation) is the key structure of this method. In this study, to assess the effect of microbial interactions on the cultivation of environmental microorganisms, soil and activated sludge samples were cultured, and colony formation and diversity during cultivation in GMDs were investigated. Results showed that the cultivation efficiency of GMD-agg was approximately 10 times higher than that of the method in which microbial interactions were canceled. In addition, the growth of taxonomies containing many uncultured microorganisms, such as *Verrucomicrobia*, *Planctomycetia*, *Acidobacteria*, and *Vicinimibacteria*, was observed. Furthermore, when isolated strains were co-cultured at high cell densities with either inter-or intra-species, we observed a higher cultivation efficiency than when microbial interactions were canceled. Rebuilding growth-promoting microbial interactions using isolated strains demonstrated that microbial interactions positively influence microbial growth. These findings indicate the effectiveness of this cultivation method, reveal the crucial role of microbial interactions in the growth of environmental microorganisms, and contribute to solving the issue of microbial uncultivability.

**IMPORTANCE:** The overwhelming majority of environmental microbes remain uncultured, limiting our understanding of their physiology and ecological roles. Although microbial interactions have been predicted as one of the key factors for the growth of uncultivable microbial types, the effect of these interactions on cultivability remains poorly understood. In this study, we developed a new droplet-based co-cultivation approach that promotes microbial interactions while maintaining pure cultures and enables growth tracking at the single-cell level. This method significantly improved cultivability (approximately 10 times), including the growth of taxa that are difficult to cultivate. Direct observation of microbial growth in the community using this method clearly demonstrated that microbial interactions are essential for the growth of diverse microbial types. These findings underscore the importance of microbial interactions in cultivation and offer a basis for radically expanding microbial bioresources, manipulating microbial communities, and exploring previously unrecognized microbial interactions.

## INTRODUCTION

The majority of microorganisms are known to be difficult to cultivate. Colony formation on solid media, such as agar plates, which are widely used as the primary cultivation technique, remains a foundational method for isolating environmental microbes. However, a substantial discrepancy often exists between the number of microbial cells inoculated from environmental samples and the number of colonies that subsequently form on artificial media (1). This long-standing issue, known as “The Great Plate Count Anomaly” (2, 3), was recognized over a century ago (4) and remains unresolved. A similar discrepancy is observed between the vast diversity of uncultivated microbial species revealed by cultivation-independent methods, such as 16S rRNA gene analysis and single-cell or meta-genomics, and the comparatively limited number of cultivated representatives (5–10). This limitation poses a major bottleneck in advancing our understanding of microbial physiology, ecology, and their potential applications. Addressing this issue requires the development of novel cultivation strategies and a deeper understanding of why many microorganisms resist cultivation under laboratory conditions.

In environments with high cell densities, such as soil, interactions among microbes are thought to regulate both growth and community structure (11–15). Indeed, some microorganisms depend on microbial interactions for growth, relying on compounds supplied by “helper” organisms (16–20). However, standard isolation procedures are designed to obtain pure cultures and typically prevent these interactions. For instance, only a limited number of cells (up to 10⁴ cells per plate) can be inoculated on agar plates to maintain sufficient spacing, thereby preventing colony overlap and cross-contamination. This separation disrupts potential interactions, especially those mediated by diffusible metabolites, eliminating many growth-supporting relationships (21). Consequently, microorganisms that require such interactions have largely been overlooked by conventional cultivation methods, except in rare or accidental cases. Consequently, the true extent to which microbial growth depends on these interactions remains largely unclear.

In contrast, *in situ* cultivation techniques, where microorganisms are grown within their natural environment using membrane-bound devices, have enabled the recovery of previously uncultivated taxa (22–27). These methods aim to incorporate environmental growth factors that are absent in standard artificial media. While some of these factors likely originate from microbial neighbors, they also include other abiotic components. The specific contribution of microbial interactions, particularly via exchanged chemical signals or metabolites, to cultivability remains poorly understood, as isolating these variables experimentally is technically challenging. Traditional cultivation methods are not suited to maintaining the close spatial proximity needed for such interactions to occur via diffusion (28–30). However, increasing both inoculum density and spatial proximity conflicts to obtain pure cultures, distinguishable colonies, creating a fundamental limitation in conventional methods.

In this study, we developed a new microbial cultivation platform using microdroplet technology. The key structure of this approach is hydrogel particles, gel microdroplets (GMDs) measuring 10–30 µm in diameter, which encapsulate single cells with a medium softly aggregated within oil. This system supports extremely high cell densities (>10⁷–10⁸ cells/mL), comparable to those in soil (31), while still preserving isolation of individual cells or micro-colonies. This enables microbial interactions to occur at spatial scales similar to those in natural environments, particularly in soils and biofilms. Crucially, the droplet-based confinement allows for the separate cultivation and tracking of individual colonies without risk of cross-contamination. We applied this method to environmental samples from soil and activated sludge to assess the influence of microbial interactions on cultivation efficiency. Furthermore, we analyzed the isolates recovered using this platform and demonstrated that both intra– and inter-species interactions significantly contributed to their growth.

## RESULTS

### Conception of the new cultivation method for high microbial interaction

We hypothesized that cultivating microorganisms at much higher cell densities than those typically used in agar plating would promote interactions through diffusible compounds, including growth factors. This, in turn, could enable the recovery of a wider range of microbial taxa compared to traditional cultivation methods. To test this hypothesis, we developed the new cultivation approach, as illustrated in Fig. 1a. This method, referred to as “Gel Microdroplet Aggregate in Oil” (GMD-agg), achieves extremely high inoculum cell densities (>10⁷–10⁸ cells/mL).

**Fig 1.**
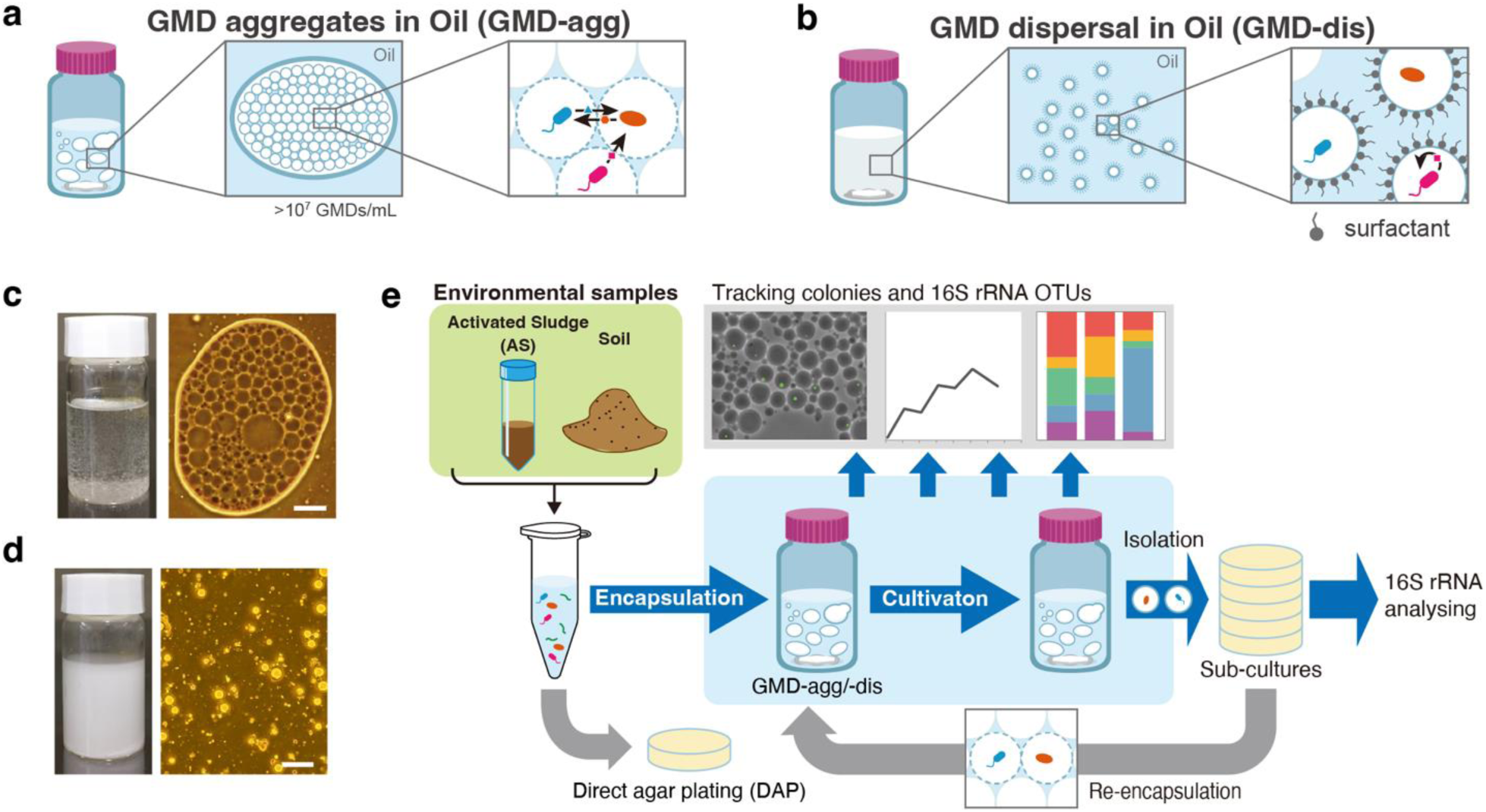
Conceptual overview of the GMD-agg cultivation method and experimental design. (a) Schematic representation of the gel microdroplet aggregation (GMD-agg) cultivation method. In this system, a large number (10⁷–10⁸) of gel microdroplets (10–30 μm in diameter), each containing a single microbial cell, are softly aggregated in oil. This structure enables the parallel cultivation of spatially separated microcultures, allowing water-soluble compounds to diffuse between droplets and facilitate microbial interactions while maintaining physical isolation between cultures. (b) Schematic representation of the gel microdroplet dispersion (GMD-dis) cultivation method. In contrast to GMD-agg, a stable emulsion is maintained, and droplets remain uniformly dispersed. Each droplet functions as a closed and independent culture unit. (c) Left: macroscopic view of the GMD-agg cultivation. Right: phase-contrast microscopic image of GMD-agg showing aggregated droplets in mineral oil. Scale bar, 50 μm. (d) Left: macroscopic view of the GMD-dis cultivation. Right: phase-contrast microscopic image of GMD-dis showing dispersed droplets in mineral oil containing 1% Span80. Scale bar, 50 μm. (e) Overview of the experimental demonstration of the new cultivation method (GMD-agg). Details are provided in the method.

GMDs were prepared by emulsifying a hydrogel (agarose) containing both a medium and suspended microbial cells in oil with a surfactant. The resulting GMDs were then transferred to mineral oil without surfactants. In the absence of surfactants, hydrophilic GMDs aggregate within the oil phase. To maintain uniform oxygen availability (aerobic conditions) and promote consistent interactions, the GMD-oil mixture was gently stirred throughout the incubation period. This produced small GMD-aggregates approximately 1–10 mm in diameter (Fig. 1c), which repeatedly came into contact with and separated from each other during cultivation. In contrast, conventional droplet-based cultivation methods maintain each droplet in a fully emulsified and dispersed state (32–34), thereby limiting chemical diffusion between droplets (Fig. 1b, d).

A key feature of the GMD-agg method is that cells can exchange water-soluble compounds through diffusion without cross-contamination while maintaining a high cell density. We demonstrated this using *Escherichia coli* as a model organism. GMDs containing *E. coli* were mixed with empty GMDs and cultured under GMD-agg conditions. After 375 hours of incubation, none of the 1,000 initially cell-free GMDs contained growing cells (Supplementary Fig. S1), confirming that colony separation was maintained and that there was no cross-contamination. We also tested the diffusion of water-soluble compounds between GMDs. GMDs containing *E. coli* were mixed with GMDs containing sodium azide, a known growth inhibitor of *E. coli* (35), and cultivated under GMD-agg conditions. After 20 hours, growth inhibition was observed in *E. coli* containing GMDs, indicating that sodium azide had diffused between droplets (Supplementary Fig. S2). These findings confirm that the GMD-agg method allows both spatial separation of growing colonies and the diffusion of soluble compounds across droplets.

In this study, we evaluated the cultivation efficiency of GMD-agg in comparison with two other methods: GMD cultivation without aggregation (GMD-dis) and conventional direct agar plating (DAP). Specifically, we focused on (1) the colony formation rate by tracking visible colonies, (2) microbial diversity and its changes during incubation, and (3) the isolation of previously uncultivated microorganisms through subculturing from GMDs to agar plates (Fig. 1e).

### GMD-agg improves the cultivation efficiency for environmental samples

#### Improving the colony formation ratio

To investigate the effect of the GMD-agg method on soil and activated sludge (AS) samples, we measured the colony formation ratio during incubation. For comparison, the same samples were cultured using the GMD-dis and DAP methods. Single microbial cells from environmental samples were encapsulated in GMDs containing 0.1× R2A medium. Cells were inoculated into GMDs following the Poisson distribution, resulting in approximately 8–12% of GMDs containing a single microbial cell.

During incubation, the colony formation ratio increased over time in the GMD-agg cultures for both soil– and AS-derived cells. In contrast, little or no increase was observed in the GMD-dis and DAP cultures, and the colony formation ratio remained low throughout the incubation period. The average maximum colony formation ratios for GMD-agg, GMD-dis, and DAP were 12.3 ± 4.6%, 1.4 ± 0.4%, and 1.4 ± 0.8%, respectively, for soil samples (Fig. 2a), and 24.6 ± 11.5%, 2.8 ± 0.9%, and 1.8 ± 0.9%, respectively, for AS samples (Fig. 2b). Multiple independent experiments using soil from the same sampling sites yielded a maximum colony formation ratio of 68% in GMD-agg cultures (Supplementary Fig. S3).

**Fig 2.**
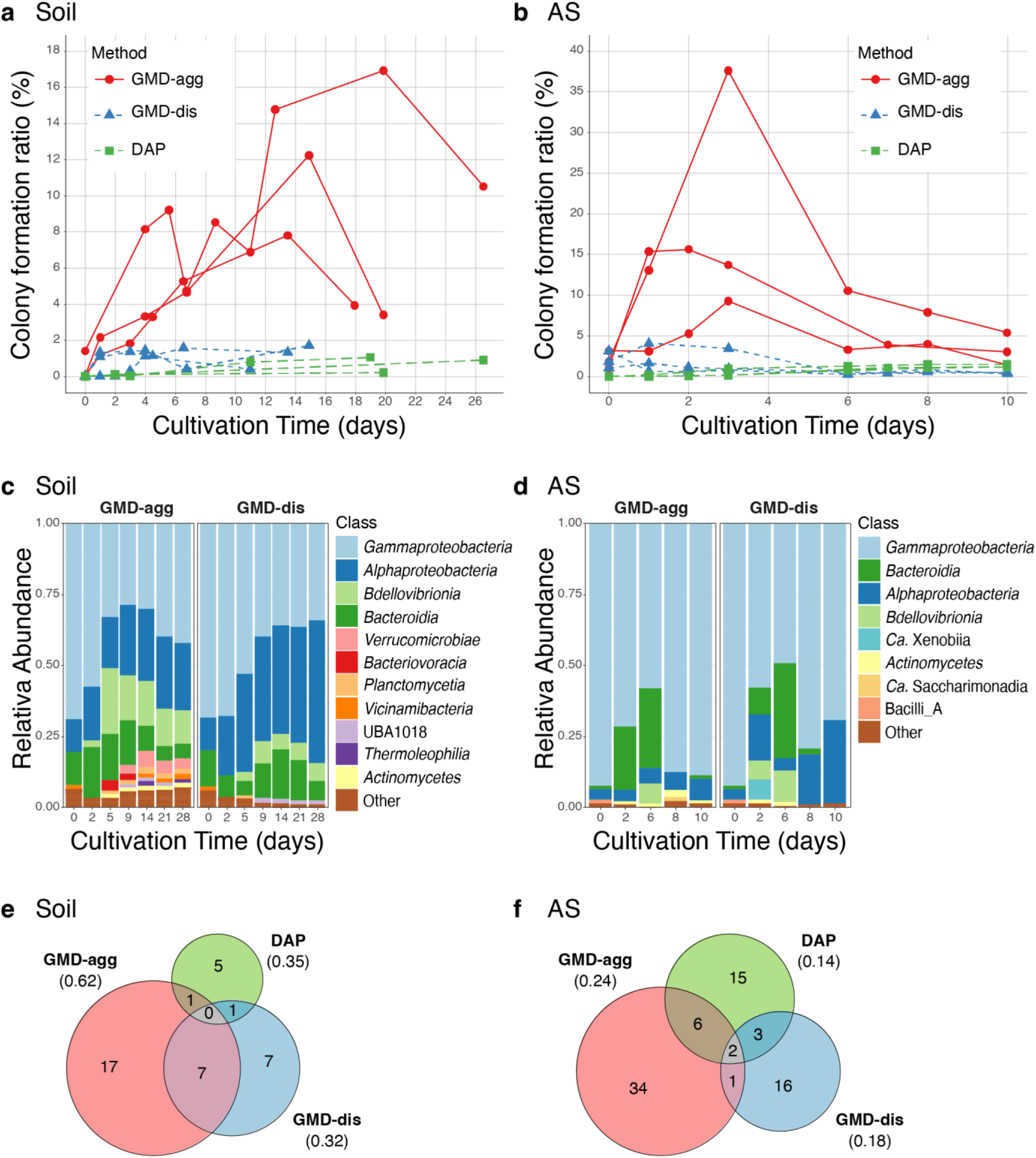
Comparison of cultivability and taxonomic composition under GMD-agg and GMD-dis cultivation conditions. (a, b) Tracking of colony formation over time in soil (a) and activated sludge (b) samples. Red circles and lines represent GMD aggregation in oil (GMD-agg), blue triangles and lines represent GMD dispersion in oil (GMD-dis), and green squares represent standard agar plate cultivation (DAP). “AS” indicates activated sludge. (c, d) Microbial community dynamics during the cultivation of soil (c) and activated sludge (d) samples. The results of GMD-agg are shown on the left and those of GMD-dis are shown on the right. ASVs with a relative abundance below 1% were grouped as “Other.” (e, f) Venn diagrams showing the number of species uniquely or commonly isolated by each method (GMD-agg, GMD-dis, and DAP) from soil (e) and activated sludge (f) samples. The numbers in parentheses indicate the proportion of species relative to the total number of isolates for each method (see Supplementary Tables S1 and S2).

These results indicate that cultivation at higher inoculum densities in GMD-agg promotes the colony formation of environmental microorganisms more effectively than DAP or GMD-dis. The similar performance of GMD-dis and DAP suggests that the increase in colony formation in GMD-agg is not simply due to encapsulation in GMDs or due to a difference in observed colony size between those in GMDs and on plates.

#### Transition of microbial community structure

Next, we examined the microbial community composition to assess how GMD-agg influenced the taxonomic structure using amplicon sequencing of the 16S rRNA V4 region. In soil samples, community structures differed markedly between GMD-agg and GMD-dis (Fig. 2c and Supplementary Fig. S4). In GMD-dis, the major taxa, including *Gammaproteobacteria*, *Alphaproteobacteria*, *Bdellovibrionia*, and *Bacteroidia*, were consistent between the early (day 2 and day 5) and late (day 21 and day 28) stages (Fig. 2c). In GMD-agg, however, while similar taxa dominated in the early period, the late-stage communities included additional taxa, such as *Verrucomicrobia*, *Actinomycetia*, *Planctomycetia*, and *Vicinimibacteria*. Notably, *Verrucomicrobia*, *Planctomycetia*, and *Vicinimibacteria* are classes with few cultivated representatives. In contrast, the AS samples showed less pronounced differences between GMD-agg and GMD-dis (Fig. 2d and Supplementary Fig. S5).

Alpha diversity analysis revealed the impact of GMD-agg on microbial diversity (Table 1). In soil, diversity indices (Faith’s PD, Shannon entropy, observed features, and Chao1) were similar between methods on day 2 but were consistently higher in GMD-agg by day 28. Particularly large differences were observed in the observed features and Chao1, which reflect the number of detected ASVs. In the AS samples, all diversity measures were also higher in GMD-agg by day 10, although the differences were smaller.

**Table 1.** α-Diversity of each GMD sample.

| Sample | Soil |  |  |  |  |  |  | AS |  |  |  |  |  |  |
| --- | --- | --- | --- | --- | --- | --- | --- | --- | --- | --- | --- | --- | --- | --- |
| Method | Source | GMD-agg |  |  | GMD-dis |  |  | Source | GMD-agg |  |  | GMD-dis |  |  |
| Time (day) | 0 | 2 | 14 | 28 | 2 | 14 | 28 | 0 | 2 | 6 | 10 | 2 | 6 | 10 |
| Faith pd | 11.0 | 7.37 | 19.1 | 18.7 | 7.99 | 7.85 | 7.75 | 12.4 | 5.39 | 6.7 | 11.5 | 11.03 | 6.9 | 10.53 |
| Shannon entropy | 6.11 | 6.38 | 7.68 | 7.64 | 6.46 | 6.80 | 7.07 | 4.53 | 5.16 | 4.65 | 3.70 | 5.48 | 5.07 | 3.68 |
| Observed features | 164 | 148 | 311 | 320 | 149 | 160 | 183 | 117 | 131 | 158 | 151 | 221 | 170 | 128 |
| Chao1 | 164 | 150 | 322 | 327 | 151 | 162 | 183 | 118 | 134 | 161 | 174 | 234 | 182 | 164 |

Beta diversity analysis based on Bray-Curtis distances further confirmed these results. Principal coordinate analysis (PCoA) showed distinct separation of the GMD-agg and GMD-dis communities in the soil throughout the incubation period (Supplementary Fig. S6a). In contrast, no major differences were observed in the AS communities between the two methods (Supplementary Fig. S6b).

#### Sub-cultivation and isolation

To compare the diversity of sub-cultivable microorganisms obtained by each method, we isolated colonies from GMD-agg, GMD-dis, and DAP cultures. Colonies from GMDs were recovered by plating them on agar and isolating individual colonies. In total, 282 isolates (94 per method) from soil and 240 isolates (80 per method) from AS were randomly selected and identified using 16S rRNA gene sequencing.

From the soil samples, 86 (GMD-agg), 77 (GMD-dis), and 90 (DAP) isolates were successfully identified, representing 21, 14, and 13 OTUs, respectively (Supplementary Table S1). For AS samples, 69 (GMD-agg), 66 (GMD-dis), and 71 (DAP) isolates were identified, corresponding to 43, 21, and 25 OTUs, respectively (Table S2). The OTU-to-isolate ratios were 0.24 (GMD-agg), 0.18 (GMD-dis), and 0.14 (DAP) for soil (Fig. 2e), and 0.62, 0.32, and 0.35 for AS (Fig. 2f), respectively. These results indicate that GMD-agg has greater taxonomic diversity among isolates in both sample types.

Moreover, the proportion of potentially novel species—defined as strains with ≤97% 16S rRNA similarity to the closest known relative—was higher among GMD-agg isolates, particularly in the AS sample (Supplementary Table S2).

#### Rebuilding growth-promoting microbial interactions using isolated strains

We next examined whether the GMD-agg cultivation method improves cultivation efficiency not only for environmental samples, but also when using isolates obtained from GMD-agg. In this experiment, we focused on both inter– and intra-species microbial interactions, specifically by testing co-cultures with helper strains and with environmental microbial communities.

To this end, we first selected strains with low colony formation ratios, isolated through GMD-agg, as test strains (Fig. S7). We then selected helper strains that promoted the colony formation of these test strains (Tables S5 and S6). Four test strains each were selected from the soil and AS. The experimental design is illustrated in Fig. 3. To evaluate recovery from a low-activity state, test strains were cultured under five different conditions: (1) DAP (pure culture), (2) GMD-dis (pure culture), (3) GMD-agg (pure culture), (4) GMD-agg co-culture with helper strains, and (5) GMD-agg co-culture with environmental microbial communities (soil or AS).

**Fig 3.**
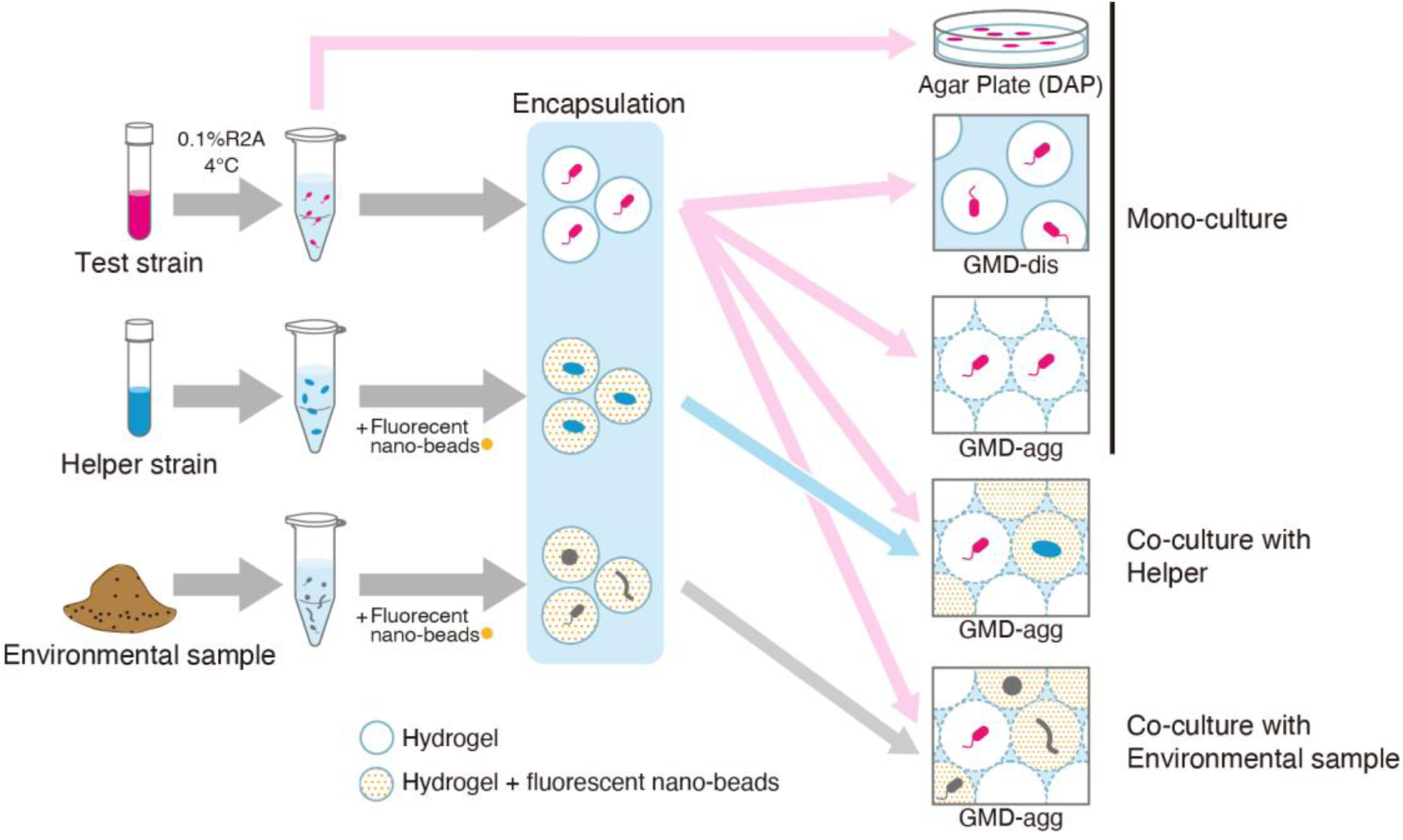
Experimental design for rebuilding growth-promoting microbial interactions using isolated strains. The test strains were first incubated at 4°C in 0.1% R2A medium to induce a low-activity state. To distinguish the droplets containing the test strains from those containing the helper strain (blue) or environmental microorganisms (gray), the droplets were fluorescently labeled by encapsulating the fluorescent nano-beads. This labeling enabled tracking of test strain growth independently under co-culture conditions. Droplets containing each cell type were prepared separately and used for co-culture in the GMD-agg system or for monoculture.

To distinguish the test strains from helper strains or environmental microbes, GMDs containing the test strains were labeled with fluorescent nanoparticles, allowing us to track their growth even under co-culture conditions (Fig. 3).

Figures 4a and 4b show the results for the soil– and AS-derived test strains, respectively. For both groups, the recovery rate (colony formation ratio) of the test strains was generally low, often below 5%, in DAP and GMD-dis cultures (except for G10 and G17 in GMD-dis). In contrast, GMD-agg co-cultured with helper strains or environmental communities often led to much higher recovery rates than those observed in pure culture systems (DAP and GMD-dis) (Fig. 4a, b).

**Fig 4.**
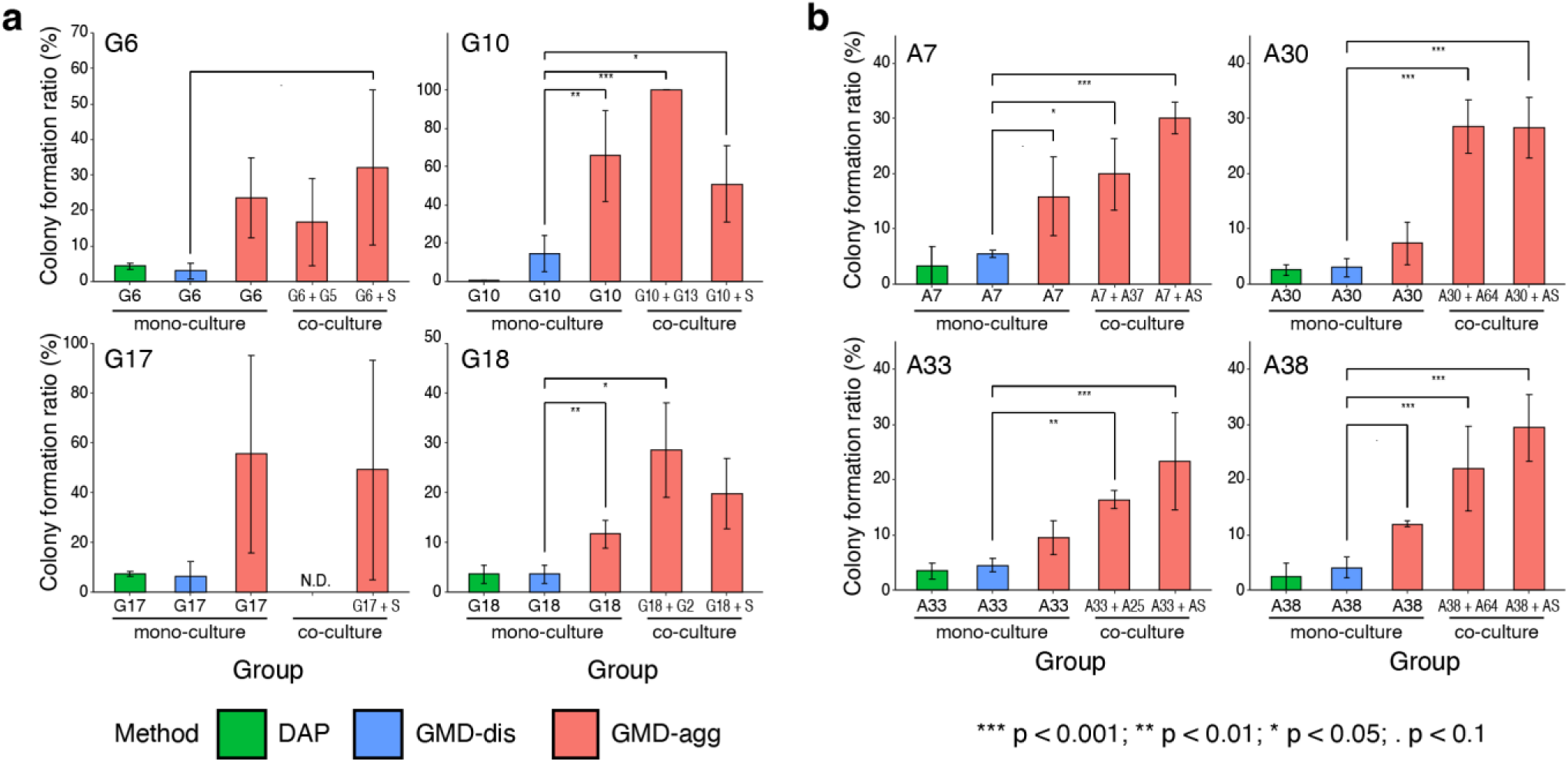
Colony formation ratios at the rebuilt growth promoting microbial interactions using isolated strains. (a) Colony formation ratios of test strains isolated from soil (G6, G10, G17, and G18) and (b) test strains isolated from AS (A7, A30, A33, and A38) under various culture conditions. “S” indicates soil samples. “AS” indicates activated sludge samples. Error bars represent the standard deviation (SD). N.D., no data.

For the soil-derived isolates, co-culture of test strains G6, G10, and G18 with their respective helper strains (G5, G13, and G2) in GMD-agg resulted in recovery rates of 17%, 100%, and 29%, respectively, which was approximately 6.7– and 8.0-fold higher than that of DAP and GMD-dis for G10 + G13 and G18 + G2. Similarly, the co-culture of G6, G10, G17, and G18 with the soil microbial community yielded recovery rates ranging from 20% to 51%, corresponding to 3.4- to 11-fold increases compared to DAP or GMD-dis (e.g., G6 + S, G10 + S) (Fig. 4a).

For the AS-derived isolates, co-culture of A7, A30, A33, and A38 with their helper strains (A3, A64, A25, and A64) in GMD-agg culture led to recovery rates that were 3.6-, 9.7-, 3.6-, and 5.4-fold higher, respectively, than in DAP and GMD-dis (e.g., A7 + A3, A30 + A64). Co-culture of these test strains with the AS microbial community also improved the recovery rates by 5.5-, 9.3-, 5.1-, and 7.0-fold, respectively (Fig. 4b).

Furthermore, even in the absence of helper strains or environmental microbes, the recovery rate of some test strains (A7, A38, G10, and G18) was 2.9- to 4.4-fold higher in GMD-agg than in DAP or GMD-dis. These results indicate that the GMD-agg method supports the recovery of low-activity cells, likely by enabling both intra– and inter-species microbial interactions.

## DISCUSSION

The significance of microbial interactions in natural environments remains poorly understood, largely due to technical limitations in assessing microbial growth at the single-cell level under high cell density conditions (>10⁷ cells/mL). In this study, we developed a new cultivation platform, termed the GMD-agg method, in which hydrogel microdroplets (GMDs) containing individual microbial cells are cultured in oil, allowing gentle aggregation. Using this system, we demonstrated that both inter– and intra-species microbial interactions are essential for the growth of diverse microbial taxa, and that such interactions are a major factor contributing to microbial uncultivability. Moreover, we successfully reconstructed growth-promoting interactions using GMD-agg–derived isolates, demonstrating that microbial interactions directly promote microbial growth.

Specifically, the colony formation ratio in GMD-agg cultures was approximately 10 times higher than that in direct agar plating (DAP). The low colony formation ratios observed in DAP (1–2% for soil and 3– 5% for AS) were consistent with previous studies (1, 2). In contrast, the GMD-dis condition, in which GMDs were physically separated, showed colony formation ratios similar to those of the DAP condition. Notably, the colony formation ratios measured by counting visible colonies on agar plates and microcolonies under a microscope in GMD-dis were comparable, indicating that the enhanced efficiency in GMD-agg is not simply due to droplet encapsulation but rather due to GMD aggregation. In GMD-agg cultures, GMDs remain loosely aggregated, maintaining a spatial distance between cells of approximately 100 μm, thereby allowing an extremely high microbial cell density (10⁸–10⁹ cells/mL) from the start of cultivation. This density approximates that found in soil (31) and is 1,000–10,000 times higher than that in standard agar plating. Under such conditions, water-soluble compounds can diffuse among GMDs, enabling microbial interactions and likely accounting for the higher cultivation efficiency compared to GMD-dis and DAP.

This observation appears inconsistent with previous reports, suggesting an inverse relationship between inoculum density and culturability (36, 37). However, other studies have reported that colony-forming units (CFUs), particularly in the early stages of incubation, can peak at specific inoculum densities (e.g., 3.94×10⁴ cells/plate), with lower densities leading to reduced CFUs (38). Thus, the relationship between inoculum density and culturability warrants careful consideration.

GMD-agg also supported the growth of a broader range of microbial taxa than GMD-dis, particularly in soil samples. This suggests that microbial interactions contribute to maintaining high microbial diversity in the soil. While some studies have reported that microbial interactions are predominantly competitive(39, 40), these findings are based on cultured strains and may not reflect the dynamics of natural microbial communities. Our results imply that positive interactions may be more prevalent in diverse uncultured communities. This view is supported by recent studies showing that high community connectivity helps to maintain microbial diversity (11).

For soil microbes, it has been reported that colony appearance timing is influenced by lag time, and that cells can be grouped into clusters based on this timing (38, 41). If lag time alone determined colony formation, GMD-dis and GMD-agg cultures would yield similar results. However, the distinct community changes observed in GMD-agg suggest that microbial interactions may reduce lag time or induce sequential growth through interaction cascades (42). Thus, microbial interactions appear to support the growth of dormant or slow-growing taxa, enabling the cultivation of a broader range of environmental microbes.

To further examine the role of microbial interactions in growth promotion, we tracked the behavior of specific isolates obtained from GMD-agg cultures. When re-inoculated into the original environmental microbial community, the recovery of the test strains increased (Fig. 4), indicating that growth-promoting interactions occur within natural communities. Although this study tested a limited number of strains, similar effects are likely to be applicable to a broader range of microbes. Recovery rates were also enhanced when the test strains were co-cultured with helper strains (Fig. 4). Additionally, for some test strains, recovery was improved even under monoculture in GMD-agg, compared to GMD-dis and DAP where microbial interactions are limited. These results suggest that both inter– and intra-species interactions contribute to microbial growth in natural settings. Therefore, initiating cultures at a high cell density, as achieved in GMD-agg, may be critical for cultivating environmental microorganisms, including previously uncultivated taxa.

However, the specificity of helper strains responsible for inter-species interactions was not fully explored in this study and warrants further investigation. It also remains unclear which substances are responsible for inducing or promoting growth and whether they trigger the initiation of proliferation or accelerate its progression. These aspects will need to be clarified in future studies.

This study introduced two methodological advances that enabled the discovery of previously unrecognized growth-inducing interactions in environmental microbial communities:

First, before this work, no method existed that could achieve high inoculum densities (i.e., intercellular distances <10 µm) similar to those in natural environments while maintaining separation between cultures and allowing the exchange of soluble factors. We accomplished this by aggregating the GMDs in oil. During cultivation, individual cultures remained isolated (i.e., pure) but were capable of interacting via diffusible compounds.

Second, by encapsulating fluorescent particles within droplets, we were able to identify and track specific cells in co-culture or complex communities. While fluorescent protein labeling is widely used, it is limited to genetically tractable strains. In contrast, our approach allows strain-specific tracking in high-density (10⁷–10⁸ cells/mL) systems that are representative of natural conditions, such as soil. Using this method, we identified various interaction patterns within microbial communities, between species, and even within species that promoted microbial growth.

## CONCLUSION

In this study, we developed a new cultivation method that promotes microbial interactions by increasing the bacterial cell density to levels comparable to those in natural environments. This approach led to a substantial increase in the colony formation of environmental microorganisms. Compared with the dispersed culturing system, the number of colonies formed increased by nearly tenfold. From a diversity standpoint, the method also maintained higher taxonomic diversity among soil bacteria than did the dispersed system. Additionally, its effectiveness was confirmed using isolated strains, further supporting the role of microbial interactions in microbial growth.

The GMD-agg cultivation method provides an effective platform for isolating and cultivating microorganisms whose growth is promoted by high inoculum density or by interactions with other microbes. Because many microorganisms in the environment are thought to grow through such interactions, this method offers a new approach to explore the ecology of environmental microbes by enabling their cultivation under interaction-permissive conditions.

## MATERIALS AND METHODS

### Environmental sample preparation

Soil and activated sludge (AS) samples were used for microbial cultivation. Soil samples were collected from a forest area located at Hiroshima University, Hiroshima, Japan. Activated sludge samples were obtained from the wastewater treatment system at the waste treatment plant in Higashihiroshima, Hiroshima, Japan. Soil samples (10 g in 100 mL of 0.1x PBS) and activated sludge samples (20 mL) were homogenized at 30% power for 1 min using an ultrasonicator. The debris in the suspensions was removed by filtration through a 30 μm-pore-size nylon net filter (Merck, USA), a 10 μm-pore-size nylon net filter, and a 5 μm-pore-size cellulose nitrate filter (25-mm diameter; Merck, USA). Microbial cells were concentrated by centrifugation at 20,000 × g at room temperature (25°C) for 10 min. For activated sludge, a subsample was additionally filtered at least three times onto a 2 μm-pore-size cellulose nitrate filter (25-mm diameter; Merck, USA) to remove initial micro-colonies. Bacterial cells were diluted in buffer (0.1x PBS, pH 7.2) to a final concentration of 10^7-8^ cells/mL.

### Encapsulation into GMD

Microbial cells (100 μL, 10^7-8^cells/ml) were mixed with 40°C preheated 1.5% low-melting agarose (0.5 mL, Seq plaque, Lonza, USA), 10% (final concentration) R2A (50 μL, Nihon Seiyaku, Japan), and 10% PLORONIC (25 μL, Sigma-Aldrich, USA). The mixture was added to 15 ml of 1% Span 80, including mineral oil (40°C, Sigma-Aldrich, USA) and emulsified at room temperature (25°C) using a CellSys 100 micro drop maker (1,500 rpm, 2 min, One Cell System). The mixture was then cooled with ice under stirring for 7 min at 150 rpm.

### Formation of GMD aggregates in oil

The solution, including the encapsulated cell, was mixed homogeneously and centrifuged at 500 × g for 5 min. To remove the surfactant, only the pellet was washed by centrifugation (500 × g for 5 min) with 10% R2A twice, and then the pure pellet containing the encapsulated cells was harvested. The retrieved microcapsules were added to 15 ml of mineral oil without surfactant.

### Incubation of GMDs in oil

The prepared samples were incubated under GMD-agg conditions at room temperature (25°C) for 10–28 days, depending on the environmental samples. To compare bacterial cultivability, the GMD-dis condition in 1% span 80 and the conventional agar plating (5,000 cells/plate) method were employed simultaneously.

### Monitoring the colony formation ratio

A subsample (600 μL) was collected by centrifugation at 500 × g at room temperature (25°C) for 5 min. The collected GMDs were washed with 1 mL MilliQ aliquots by centrifugation at 500 × g for 5 min in triplicate. The cells in the microdroplet were stained with SYBR Green solution (10^-3^ Molecular probes, USA) for 5 min in the dark. The number of microcolonies(cell to microcolony) was monitored throughout the incubation period. On the initial day (0 h), the single cell number in GMD was counted to calculate the cultivability. In the conventional agar plating method, colony-forming units are monitored during cultivation.

The colony formation ratio of in-oil cultivation (GMD-agg and GMD-dis) was represented as the microcolony number in the microcapsule to the total inoculum density and calculated using equation (1). Agar plating (colony formation ratio of DAP) was estimated using equation (2). where t is the cultivation time, M is the total number of micro-droplets, and G and C are the total number of gel micro-droplets and single cells, respectively. CFU is the colony-forming unit, and I is the inoculum size as a constant.

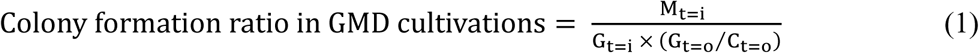

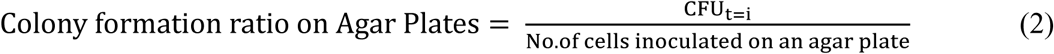

### Community diversity analysis

Community diversity was analyzed using amplicon sequencing targeting the V4 region of 16S rRNA. Microbial DNA was extracted from the GMD after washing using the DNeasy Soil Pro kit (QIAGEN, Netherlands) according to the manufacturer’s instruction and then stored at –28°C.

DNAs obtained from soil samples were targeted to the V4 region of 16S rRNA, and DNAs were amplified with a 515f/805r primer set that included Illumina’s barcode and adaptor sequences. Equal amounts of amplified DNA from each sample were pooled and pair-end sequenced on the Illumina MiSeq platform at the Bioengineering Lab. Co., Ltd. DNAs obtained from activated sludge samples were targeted to the V4 region of 16S rRNA, and DNAs were amplified with a 515f/806r primer set that included a barcode sequence for sample identification. Equal amounts of amplified DNA from each sample were pooled and single-end sequenced on the Ion PGM (Thermo Fisher) platform. Raw reads of 16S rRNA amplicon sequencing were processed according to the qiime2 pipeline, and quality filtering, paired-end read merging, and chimera checking were performed using the DADA2. Sequence classification was based on the database Greengenes2 2024.09 (43). They were then plotted using multidimensional scaling based on the Bray-Curtis distance calculated using the R phyloseq package (1.42.0) (44).

### Sub-culturing from the GMD cultivations

GMDs, including cells for sub-culturing, were collected as mentioned above to track the microcolonies. Collected GMDs were added to PercollTM PLUS/Percoll (0.45 mL, GE Healthcare, USA), 50% Tween 40 (2 μL, Nacalai Tesque, Japan), and 0.15 M NaCl (1 mL, Nacalai Tesque, Japan) and centrifuged at 20,000 × g for 1 h. Pure GMDs were retrieved and washed with 1 mL Milli-Q aliquots by centrifugation at 10,000 rpm for 5 min. Approximately 30,000 GMDs were added to 10 mL of tempered (40°C) 1% sea plaque (Lonza, USA), mixed, poured onto each 10% R2A agar plate by dividing into three equal amounts, and incubated at room temperature (25°C).

### Identification of isolates by 16S rRNA sequencing

After incubation, 80-100 colonies were randomly selected from each cultivation method. Isolates were subjected to colony direct PCR. The PCR products were sequenced at Takara Bio Inc.. (Japan). Considered clones, as sequences longer than 500 nucleotides (nt) and non-chimeric, were estimated with databases in GenBank (http://blast.ncbi.nih.gov/) (45, 46) to identify the closest matching sequences using BLASTn (47).

### Screening to select test strains of low recovery

We selected strains exclusively isolated by GMD-agg cultivation. To select a specific strain with low recovery, the isolates were incubated in 10% R2A medium until they reached the stationary phase and then placed in a low activity state by starving the cells for three days at 4°C in 0.1% R2A broth. Each starved strain was inoculated onto a 10% R2A agar plate and incubated at room temperature (25°C) for 1 week. The recovery ratio was calculated as the ratio of the inoculated 500 cell density to the number of colony-forming units. As a result, four strains were selected as test strains from among the strains with a recovery rate between 0.5 and 10% (Supplementary Fig. S7).

### Screening the strains as helper strains for the co-cultivation with the test strain

To track the recovery of a test strain during co-culturing, we selected a helper strain. To select helper strains, culture supernatants were extracted from the cultures of isolates exclusively isolated by GMD-agg cultivation and incubated with 10% R2A broth. The supernatants were then autoclaved. Finally, 10% R2A agar plates containing each culture supernatant (final concentration 1%) were prepared. The 500 cells of test strains after starvation were inoculated on the agar plate with the modified 10% R2A (including supernatants) and the plate with standard 10% R2A, and the recovery ratio was calculated and compared with the ratio from the standard 10% R2A plate (Supplementary Table S5 and S6). Finally, one helper strains were selected for each strain from the activated sludge and 0 or 1 for each strain from the soil. The helper strains were used for the following co-cultivation test.

### Tracking the recovery of the test strain in co-culturing oil cultivation

To track the recovery of test strains, the test strain must be distinguished from other microbial types in co-culture with the helper strain or co-culture with the microbial community during incubation. For this purpose, the GMD was fluorescently labeled using fluorescent nano-beads. Approximately 10–30 fluorescent nano-beads (Polyscience, Inc., USA) were encapsulated in each GMD by simply mixing fluorescent nano-beads with microbial cell samples and the gelling agent when making GMDs. In the case of co-culturing with a specific helper strain, the test strain cells were encapsulated in non-fluorescent labeled GMDs, whereas the helper strain cells were encapsulated in a fluorescent labelled gel, including fluorescent nano-beads. After preparing the two types of GMDs (GMDS with test strain and GMDs with helper strain or the environmental microbial community) separately, each GMD was mixed homogeneously and incubated in oil as a co-culture for 240 h, according to the procedure of GMD-agg cultivation described above. Simultaneously, the growth of the test strains was examined in GMD-dis culture and standard agar plates with the same medium as the GMD-agg cultivation. The recovery rate of the test strain as a colony formation rate was compared among the three culture conditions.

## ACKNOWLEDGEMENTS

We acknowledge Dawoon Jung for helpful insight and discussions. We also thank Fumiko Irisuna for sequencing the 16S rRNA amplicons and the Bioengineering Lab. Co., Ltd. for sequencing the 16S rRNA amplicons. This work was supported by JSPS KAKENHI Grant Numbers JP26709063, JP19H02873, JP20H05587. R.S. was in part supported by JST, the establishment of university fellowships towards the creation of science technology innovation, Grant Number JPMJFS2129.

## DATA AVAILABILITY STATEMENT

Amplicon sequence data were deposited to NCBI under the BioProject accession number PRJDB20252. For isolated strains, 16S rRNA gene sequences were deposited to GenBank under the accession numbers 8860-8907, ON680767-ON680791, ON819649-ON819718, PP716552-PP716569, and PP843538.

## CONFLICTS OF INTEREST

The authors declare no conflict of interest.

## REFERENCES

1. Zheng M, Wen L, He C, Chen X, Si L, Li H, Liang Y, Zheng W, Guo F. 2024. Sequencing-guided re-estimation and promotion of cultivability for environmental bacteria. Nat Commun 15:9051.

2. Staley JT, Konopka A. 1985. Measurement of in situ activities of nonphotosynthetic microorganisms in aquatic and terrestrial habitats. Annu Rev Microbiol 39:321–346.

3. Amann RI, Ludwig W, Schleifer KH. 1995. Phylogenetic identification and in situ detection of individual microbial cells without cultivation. Microbiol Rev 59:143–169.

4. Conn HJ. 1918. The microscopic study of bacteria and fungi in soil. N Y Agric Exp Sta Tech Bull 64:3–20.

5. Steen AD, Crits-Christoph A, Carini P, DeAngelis KM, Fierer N, Lloyd KG, Cameron Thrash J. 2019. High proportions of bacteria and archaea across most biomes remain uncultured. ISME J.

6. Parks DH, Chuvochina M, Waite DW, Rinke C, Skarshewski A, Chaumeil P-A, Hugenholtz P. 2018. A standardized bacterial taxonomy based on genome phylogeny substantially revises the tree of life. Nat Biotechnol 36:996–1004.

7. Castelle CJ, Banfield JF. 2018. Major new microbial groups expand diversity and alter our understanding of the tree of life. Cell 172:1181–1197.

8. Schulz F, Eloe-Fadrosh EA, Bowers RM, Jarett J, Nielsen T, Ivanova NN, Kyrpides NC, Woyke T. 2017. Towards a balanced view of the bacterial tree of life. Microbiome 5:140.

9. Lloyd KG, Steen AD, Ladau J, Yin J, Crosby L. 2018. Phylogenetically Novel Uncultured Microbial Cells Dominate Earth Microbiomes. mSystems 3.

10. Achtman M, Wagner M. 2008. Microbial diversity and the genetic nature of microbial species. Nat Rev Microbiol 6:431–440.

11. Dubey M, Hadadi N, Pelet S, Carraro N, Johnson DR, van der Meer JR. 2021. Environmental connectivity controls diversity in soil microbial communities. Commun Biol 4:492.

12. Bickel S, Or D. 2020. Soil bacterial diversity mediated by microscale aqueous-phase processes across biomes. Nat Commun 11:116.

13. Romdhane S, Spor A, Aubert J, Bru D, Breuil M-C, Hallin S, Mounier A, Ouadah S, Tsiknia M, Philippot L. 2022. Unraveling negative biotic interactions determining soil microbial community assembly and functioning. ISME J 16:296–306.

14. Zelezniak A, Andrejev S, Ponomarova O, Mende DR, Bork P, Patil KR. 2015. Metabolic dependencies drive species co-occurrence in diverse microbial communities. Proc Natl Acad Sci U S A 112:6449–6454.

15. Pande S, Kost C. 2017. Bacterial Unculturability and the Formation of Intercellular Metabolic Networks. Trends Microbiol 25:349–361.

16. Morris J. Jeffrey, Kirkegaard Robin, Szul Martin J., Johnson Zackary I., Zinser Erik R. 2008. Facilitation of Robust Growth of *Prochlorococcus* Colonies and Dilute Liquid Cultures by “Helper” Heterotrophic Bacteria. Appl Environ Microbiol 74:4530–4534.

17. Ohno M, Shiratori H, Park MJ, Saitoh Y, Kumon Y, Yamashita N, Hirata A, Nishida H, Ueda K, Beppu T. 2000. Symbiobacterium thermophilum gen. nov., sp. nov., a symbiotic thermophile that depends on co-culture with a Bacillus strain for growth. Int J Syst Evol Microbiol 50 Pt 5:1829–1832.

18. Katayama T, Nobu MK, Imachi H, Hosogi N, Meng X-Y, Morinaga K, Yoshioka H, Takahashi HA, Kamagata Y, Tamaki H. 2024. A Marine Group A isolate relies on other growing bacteria for cell wall formation. Nat Microbiol 9:1954–1963.

19. Nichols D., Lewis K., Orjala J., Mo S., Ortenberg R., O’Connor P., Zhao C., Vouros P., Kaeberlein T., Epstein S. S. 2008. Short Peptide Induces an “Uncultivable” Microorganism To Grow In Vitro. Appl Environ Microbiol 74:4889–4897.

20. D’Onofrio A, Crawford JM, Stewart EJ, Witt K, Gavrish E, Epstein S, Clardy J, Lewis K. 2010. Siderophores from neighboring organisms promote the growth of uncultured bacteria. Chem Biol 17:254–264.

21. Alain K, Querellou J. 2009. Cultivating the uncultured: limits, advances and future challenges. Extremophiles 13:583–594.

22. Remenár M, Karelová E, Harichová J, Zámocký M, Kamlárová A, Ferianc P. 2015. Isolation of previously uncultivable bacteria from a nickel contaminated soil using a diffusion-chamber-based approach. Appl Soil Ecol 95:115–127.

23. Ferrari BC, Binnerup SJ, Gillings M. 2005. Microcolony cultivation on a soil substrate membrane system selects for previously uncultured soil bacteria. Appl Environ Microbiol 71:8714–8720.

24. Berdy B, Spoering AL, Ling LL, Epstein SS. 2017. In situ cultivation of previously uncultivable microorganisms using the ichip. Nat Protoc 12:2232–2242.

25. Nichols D., Cahoon N., Trakhtenberg E. M., Pham L., Mehta A., Belanger A., Kanigan T., Lewis K., Epstein S. S. 2010. Use of Ichip for High-Throughput In Situ Cultivation of “Uncultivable” Microbial Species. Appl Environ Microbiol 76:2445–2450.

26. Jung D, Aoi Y, Epstein SS. 2016. In situ cultivation allows for recovery of bacterial types competitive in their natural environment. Microbes Environ 31:456–459.

27. Kaeberlein T, Lewis K, Epstein SS. 2002. Isolating “uncultivable” microorganisms in pure culture in a simulated natural environment. Science 296:1127–1129.

28. van Tatenhove-Pel RJ, Rijavec T, Lapanje A, van Swam I, Zwering E, Hernandez-Valdes JA, Kuipers OP, Picioreanu C, Teusink B, Bachmann H. 2021. Microbial competition reduces metabolic interaction distances to the low µm-range. ISME J 15:688–701.

29. Dal Co A, van Vliet S, Kiviet DJ, Schlegel S, Ackermann M. 2020. Short-range interactions govern the dynamics and functions of microbial communities. Nat Ecol Evol 4:366–375.

30. Gantner S, Schmid M, Dürr C, Schuhegger R, Steidle A, Hutzler P, Langebartels C, Eberl L, Hartmann A, Dazzo FB. 2006. In situ quantitation of the spatial scale of calling distances and population density-independent N-acylhomoserine lactone-mediated communication by rhizobacteria colonized on plant roots: In situ quantitation of the spatial scale of calling distances. FEMS Microbiol Ecol 56:188–194.

31. Portillo MC, Leff JW, Lauber CL, Fierer N. 2013. Cell size distributions of soil bacterial and archaeal taxa. Appl Environ Microbiol 79:7610–7617.

32. Saito K, Ota Y, Tourlousse DM, Matsukura S, Fujitani H, Morita M, Tsuneda S, Noda N. 2021. Microdroplet-based system for culturing of environmental microorganisms using FNAP-sort. Sci Rep 11:9506.

33. Mahler L, Niehs SP, Martin K, Weber T, Scherlach K, Hertweck C, Roth M, Rosenbaum MA. 2021. Highly parallelized droplet cultivation and prioritization of antibiotic producers from natural microbial communities. Elife 10.

34. Eun Y-J, Utada AS, Copeland MF, Takeuchi S, Weibel DB. 2011. Encapsulating bacteria in agarose microparticles using microfluidics for high-throughput cell analysis and isolation. ACS Chem Biol 6:260–266.

35. Lichstein HC, Soule MH. 1944. Studies of the effect of sodium azide on microbic growth and respiration. J Bacteriol 47:221–230.

36. Davis KER, Joseph SJ, Janssen PH. 2005. Effects of growth medium, inoculum size, and incubation time on culturability and isolation of soil bacteria. Appl Environ Microbiol 71:826–834.

37. Bakken LR, Olsen RA. 1987. The relationship between cell size and viability of soil bacteria. Microb Ecol 13:103–114.

38. Ernebjerg M, Kishony R. 2012. Distinct growth strategies of soil bacteria as revealed by large-scale colony tracking. Appl Environ Microbiol 78:1345–1352.

39. Foster KR, Bell T. 2012. Competition, not cooperation, dominates interactions among culturable microbial species. Curr Biol 22:1845–1850.

40. Chang C-Y, Bajić D, Vila JCC, Estrela S, Sanchez A. 2023. Emergent coexistence in multispecies microbial communities. Science 381:343–348.

41. Hattori T, Mitsui H, Haga H, Wakao N, Shikano S, Gorlach K, Kasahara Y, el-Beltagy A, Hattori R. 1997. Advances in soil microbial ecology and the biodiversity. Antonie Van Leeuwenhoek 72:21–28.

42. Jung D, Machida K, Nakao Y, Kindaichi T, Ohashi A, Aoi Y. 2021. Triggering Growth via Growth Initiation Factors in Nature: A Putative Mechanism for in situ Cultivation of Previously Uncultivated Microorganisms. Front Microbiol 12:537194.

43. McDonald D, Jiang Y, Balaban M, Cantrell K, Zhu Q, Gonzalez A, Morton JT, Nicolaou G, Parks DH, Karst SM, Albertsen M, Hugenholtz P, DeSantis T, Song SJ, Bartko A, Havulinna AS, Jousilahti P, Cheng S, Inouye M, Niiranen T, Jain M, Salomaa V, Lahti L, Mirarab S, Knight R. 2024. Greengenes2 unifies microbial data in a single reference tree. Nat Biotechnol 42:715–718.

44. McMurdie PJ, Holmes S. 2013. phyloseq: an R package for reproducible interactive analysis and graphics of microbiome census data. PLoS One 8:e61217.

45. Sayers EW, Bolton EE, Brister JR, Canese K, Chan J, Comeau DC, Connor R, Funk K, Kelly C, Kim S, Madej T, Marchler-Bauer A, Lanczycki C, Lathrop S, Lu Z, Thibaud-Nissen F, Murphy T, Phan L, Skripchenko Y, Tse T, Wang J, Williams R, Trawick BW, Pruitt KD, Sherry ST. 2022. Database resources of the national center for biotechnology information. Nucleic Acids Res 50:D20–D26.

46. Sayers EW, Cavanaugh M, Clark K, Ostell J, Pruitt KD, Karsch-Mizrachi I. 2020. GenBank. Nucleic Acids Res 48:D84–D86.

47. Altschul SF, Gish W, Miller W, Myers EW, Lipman DJ. 1990. Basic local alignment search tool. J Mol Biol 215:403–410.

